# Clinically-relevant T cell expansion protocols activate distinct cellular metabolic programs and phenotypes

**DOI:** 10.1101/2021.08.24.457536

**Authors:** Sarah MacPherson, Sarah Keyes, Marisa Kilgour, Julian Smazynski, Jessica Sudderth, Tim Turcotte, Adria Devlieger, Jessie Yu, Kimberly S. Huggler, Jason R. Cantor, Ralph J. DeBerardinis, Christopher Siatskas, Julian J. Lum

## Abstract

*Ex vivo* expansion conditions used to generate T cells for immunotherapy are thought to adopt metabolic phenotypes that impede therapeutic efficacy *in vivo*. The comparison of five different culture media used for clinical T cell expansion revealed unique optima based on different output variables including proliferation, differentiation, function, activation and mitochondrial phenotypes. T cells adapted their metabolism to match their media expansion condition as shown by glucose and glutamine uptake, and patterns of glucose isotope labeling. However, adoption of these metabolic phenotypes was uncoupled to T cell function. Furthermore, T cell products cultured in ascites from ovarian cancer patients displayed suppressed mitochondrial activity and function irrespective of the *ex vivo* expansion media. In one case, culturing in ascites resulted in increased glucose uptake which was insufficient to rescue T cell function. Thus, *ex vivo* T cell expansion conditions have profound impacts on metabolism and function.

## INTRODUCTION

Cell-based immunotherapies are garnering considerable attention for their promise in treating human cancers. These therapies involve isolating and expanding autologous human tumor-infiltrating lymphocytes (TIL) or genetically modified chimeric antigen receptor (CAR) T cells. In particular, CAR T cell therapy has achieved exceptional response rates in hematological cancers^1,2^. However several barriers exist that impede the efficacy of cell based therapies targeting solid tumors^3–5^. These include but are not limited to tumor antigen escape, restricted intraepithelial trafficking, limited persistence and an inhospitable tumor microenvironment (TME)^6–8^. Another critical factor is the complex cell manufacturing process that can generate products with undesirable metabolic phenotypes and ultimately hamper *in vivo* functionality^9–11^. Culture methods that maintain metabolic profiles favoring central memory phenotypic subsets have been shown to highly correlate with improved clinical outcomes^12^. A recent study identified medium-dependent transcriptional responses in several metabolic pathways during early activation, which may be crucial in programming T cells to a committed phenotype^13–17^. At the present time, there has yet to be a universally accepted formulation to manufacture T cells making it a major challenge not only to cross-compare different clinical trials, but also to understand the metabolic parameters that may be responsible for the behavior of T cells post-transfer.

The first synthetic complex medium used for *in vitro* expansion of lymphocytes was RPMI 1640^18^, and is widely used in the Rapid Expansion Protocol (REP) of TILs^19^. Improvements in formulations that incorporate defined serum components have led to media formulations such as X-VIVO 15 and AIM-V. However, recent evidence has indicated that cells maintained under these non-physiological conditions activate different signaling and metabolic pathways compared to analogous cells generated *in vivo*^20^. This has led to the generation of culture media that recapitulate many physiological characteristics of human plasma such as Human Plasma-like Medium (HPLM)^21^ and others^22^. However, these formulations may not recapitulate all the nutritional environmental cues required to induce and sustain robust expansion specific to T cells necessitating their supplementation with human AB serum. In contrast, ImmunoCult™-XF T Cell Expansion Medium (ICM) is a serum-free, xeno-free medium that supports robust T cell expansion.

Here we investigated the influence of five cell culture media conditions on T cell metabolism, proliferation, differentiation, activation and function. Although each condition supported cell proliferation, the extent of expansion varied as did differentiation, function, mitochondrial phenotypes and metabolism. The condition with the greatest proliferation and function (REP in CTL:AIM-V medium) displayed a preference for glucose uptake over glutamine. This dominance for glucose uptake could be imposed as T cells adopted the preference for glucose when switched in CTL:AIM-V irrespective of the condition used for the initial expansion. However the change in metabolism did not appear to be linked to increased function as the switch to CTL:AIM-V differentially impacted TNFα and IFNγ production. Carbon tracing revealed REP expanded T cells having increased labeled lactate whereas ICM expanded T cells displayed increased labeling of one-carbon metabolism and entry into the TCA cycle. These patterns of labeling were media dependent as switching ICM expanded T cells into CTL:AIM-V medium reverted T cells back to higher lactate labeling as seen in the REP expanded T cells, which was associated with an increase in CD25+ and PD-1+ populations. All five T cell products were exposed to the ascites derived from ovarian cancer patients and experienced suppressed T cell function and mitochondrial activity that could not be overcome by increased glucose uptake alone. Thus, our studies highlight the impact of cell culture media on the metabolic programs, phenotypes and function of human T cells for potential use in immunotherapy.

## RESULTS

### Expansion conditions skew proliferation, differentiation and activation

To examine the influence of cell culture media on T cell products, we conducted a 12-day comparison of 5 expansion protocols (**Table 1**). We compared one of the earliest tumor infiltrating lymphocyte (TIL) expansion methods commonly referred to as the rapid expansion protocol (REP)^19^ to platforms by: STEMCELL (ICM), Miltenyi (TAC) and Corning (CMM). We also tested Human Plasma-like Medium (HPLM), which was recently developed specifically to recapitulate the human plasma environment *in vitro*^13^. Enriched CD3+ T cells were isolated from apheresis products of six different healthy donors and expanded over the course of 12 days following the manufacturers’ recommended protocols using their respective cell culture media (**Table 1**). Unlike the report by Leney-Greene et al.^13^, we were primarily interested in the longitudinal changes after expansion and the T cell phenotypes produced after 12-14 days, a time course that is similar to most clinically-approved schemas used in manufacturing T cells for cancer immunotherapy.

**Table 1:**
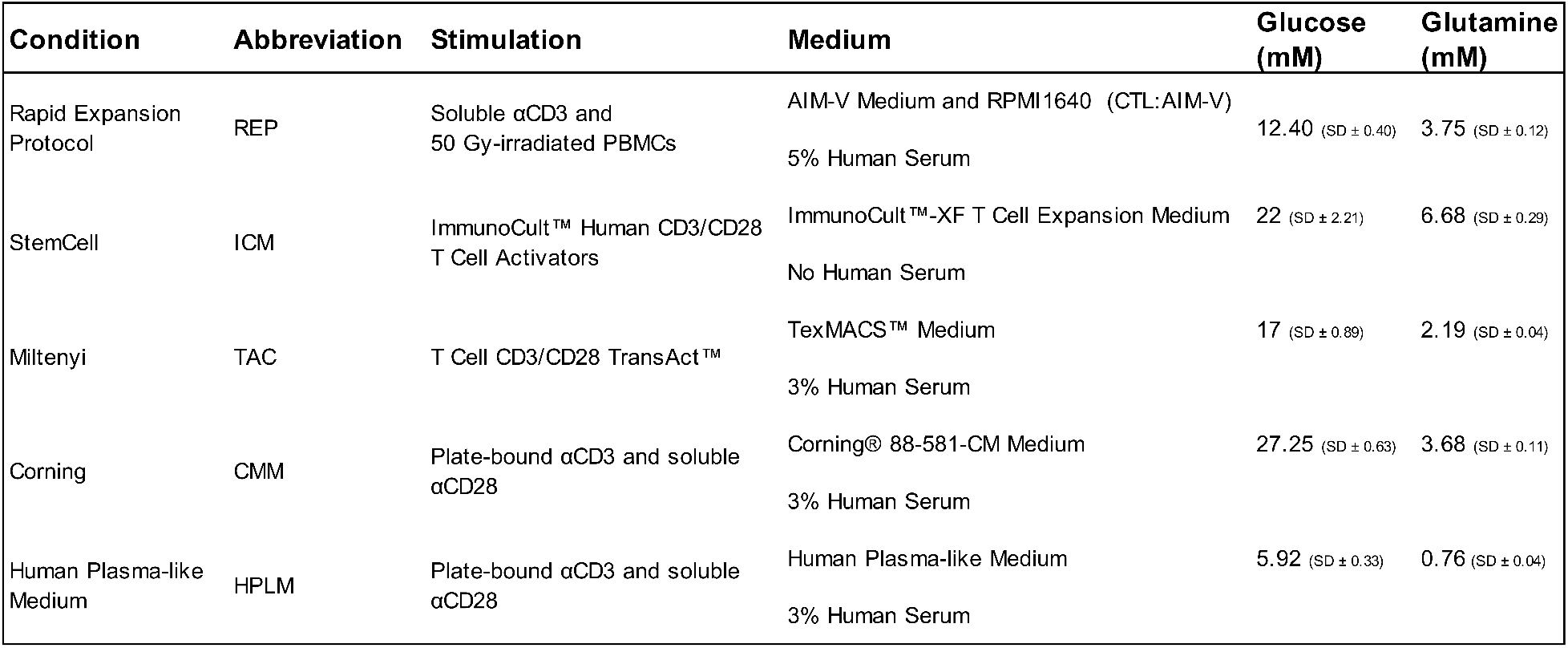
List of culture formulations and expansion conditions. Glucose and glutamine concentrations measured using colorimetric assay.

| Condition | Abbreviation | Stimulation | Medium | Glucose (mM) | Glutamine (mM) |
| --- | --- | --- | --- | --- | --- |
| Rapid Expansion Protocol | REP | Soluble αCD3 and 50 Gy-irradiated PBMCs | AIM-V Medium and RPMI1640 (CTL:AIM-V)<br>5% Human Serum | 12.40 (SD ± 0.40) | 3.75 (SD ± 0.12) |
| StemCell | ICM | ImmunoCult™ Human CD3/CD28 T Cell Activators | ImmunoCult™-XF T Cell Expansion Medium<br>No Human Serum | 22 (SD ± 2.21) | 6.68 (SD ± 0.29) |
| Miltenyi | TAC | T Cell CD3/CD28 TransAct™ | TexMACS™ Medium<br>3% Human Serum | 17 (SD ± 0.89) | 2.19 (SD ± 0.04) |
| Corning | CMM | Plate-bound αCD3 and soluble αCD28 | Corning® 88-581-CM Medium<br>3% Human Serum | 27.25 (SD ± 0.63) | 3.68 (SD ± 0.11) |
| Human Plasma-like Medium | HPLM | Plate-bound αCD3 and soluble αCD28 | Human Plasma-like Medium<br>3% Human Serum | 5.92 (SD ± 0.33) | 0.76 (SD ± 0.04) |

Given the clinical importance of maximizing the number of T cells in the expansion product, the proliferative capacity of all five of these culture medias were compared. Expansion differed significantly across all of the conditions ranging from 40-860 fold (**Fig. 1a**). This difference in expansion appeared as early as day 4 and was maintained throughout the entire expansion with the REP condition yielding the largest number of T cells. While the CMM and HPLM expansion conditions used the same stimulation method and only differed in media composition, CMM expanded cells produced a 2-fold greater expansion compared to the same T cells expanded in HPLM. It has been reported that an equivalent ratio of CD4+ and CD8+ T cells in the final infusion product is associated with better clinical responses to TIL or CAR T cell therapy^28,29^. Although each condition resulted in a variable percentage of CD4+ and CD8+ T cells, the CD4:CD8 T cell ratio within each condition was consistent (**Fig. 1b**). For example, the REP promoted a strong enrichment of CD4+ T cells and a concomitant reduction in CD8+ T cells that was significantly different than the other conditions (**Fig. 1b and Supplementary Fig. 1a**). Of note, the TAC and ICM expansion media did not appear to significantly alter the percentage of CD4+ and CD8+ T cells from day 0, whereas CMM and HPLM media generated more CD8+ T cells over the same 12-day period (**Fig. 1b**).

**Fig. 1:**
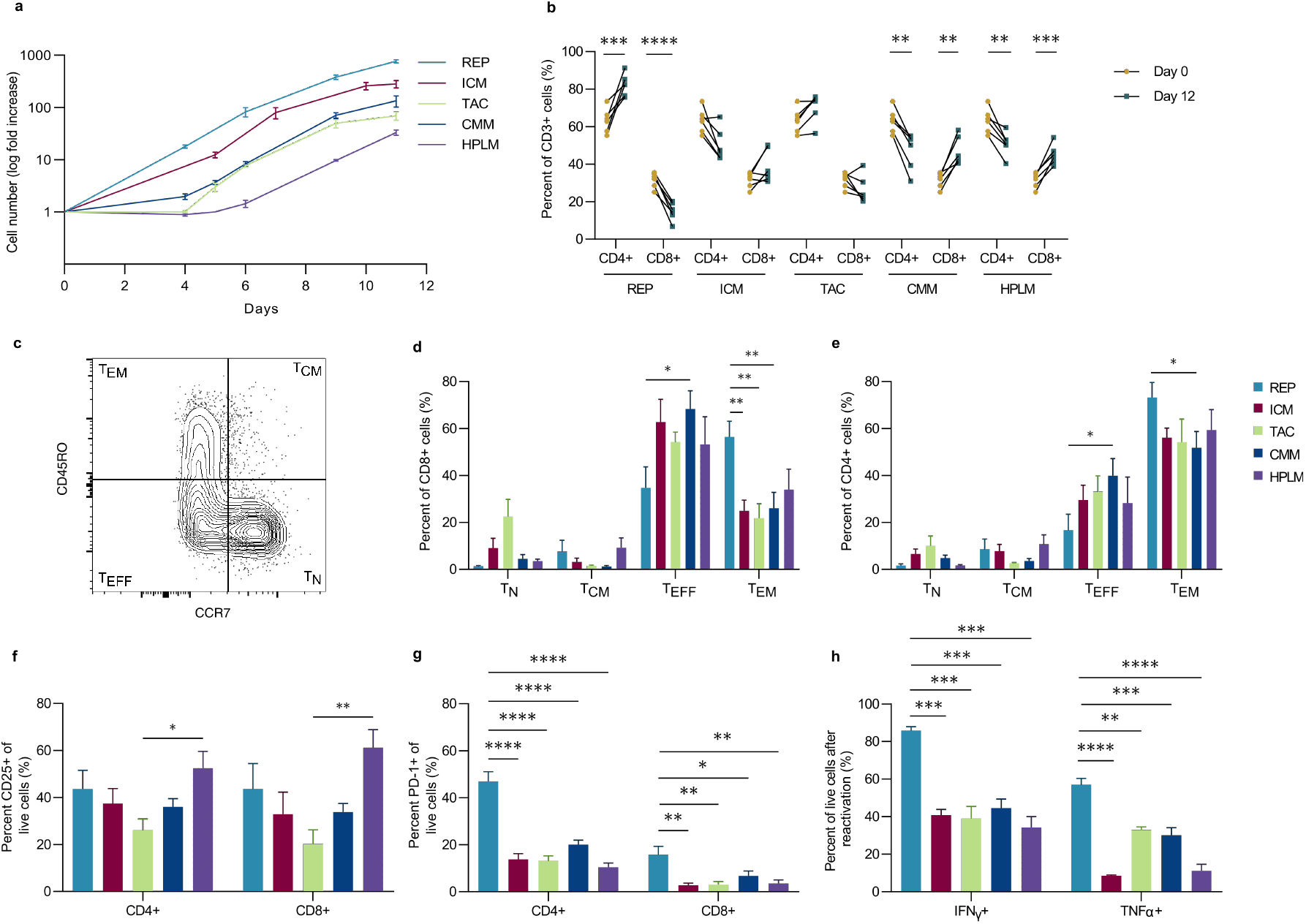
Expansion conditions skew proliferation, differentiation and activation. **(a-g)** T cells from six healthy donors were expanded in 5 different conditions for 12 days: CTL:AIM-V (REP), ImmunoCult-XF (ICM), TexMACS (TAC), Complete Corning media (CMM) and Human Plasma-like Medium (HPLM) **(a)** Log fold increase in cell number throughout the 11-day expansion **(b)** Proportion of CD4+ and CD8+ T cells of live CD3+ cells, pre- and post-expansion from six healthy donors. **(c-e)** Representative plot **(c)** and tabulated data for CD8+ **(d)** and CD4+ **(e)** T cells following expansion: naïve (T_N_), central memory (T_CM_), effector (T_EFF_), and effector memory (T_EM_). **(f)** Percentage of CD25 and **(g)** PD-1 positive CD4+ and CD8+ T cells. **(h)** Percentage of IFNγ and TNFα positive cells after CD3/CD28 reactivated following expansion (n=3). Bar graphs represent mean of n=6 **(a-g)** and **(h)** n=3 +SEM from healthy donors. Statistical significance was calculated using a Student’s t-test (**b**) or a one-way ANOVA (**d-h**) (* p<0.05, ** p<0.01, *** p<0.001, **** p<0.0001).

The differentiation state of the post-expanded population correlates with long-term persistence of both TIL and CAR T cells, and to a large extent tumor control^30,31^. For instance, the skewing towards CD8+ T effector (T_EFF_) cells limits anti-tumor responses after adoptive cell therapy^32^. On the other hand, adoptive transfer of CD8+ T central memory (T_CM_) cells has been shown to have a superior anti-tumor response than CD8+ T effector memory (T_EM_) cells^33^. Across all of the conditions tested, expanded CD4+ and CD8+ T cells displayed a predominant T_EFF_ and/or T_EM_ phenotype with the REP condition producing the highest percentage of T_EM_ cells (**Fig. 1c,d,e**). Overall, the five media conditions generated few (<10%) CD4+ or CD8+ T cells displaying a T_CM_ differentiated phenotype.

Clonal variation, marked by a diverse TCR Vβ repertoire, enables T cells to recognize a wide array of epitopes including those from tumors. Selective expansion of tumor reactive clones has been reported to correlate with clinical responses^34^. It is unclear whether certain media conditions favor the outgrowth of specific clonotypes. Therefore, we profiled TCR Vβ repertoires using flow cytometry to assess population clonality for each condition. Overall the Vβ spectratyping revealed that TCR diversity is largely maintained across all conditions (**Supplementary Fig. 1b,c)**.

To determine whether culture conditions influenced T cell activation, we measured the expression of CD25 and CD137 by flow cytometry. Expression of CD137 was generally higher on CD8+ T cells than CD4+ T cells across all conditions (**Supplementary Fig. 1d**). Although the REP produced the lowest frequency of CD8+ T cells, it achieved the highest CD8+ CD137+ cells population compared to all other conditions (**Supplementary Fig. 1d**). In contrast, CD25 expression was similar between CD4+ and CD8+ T cells, with the HPLM condition producing highest proportion of CD25+ cells compared to other conditions (**Fig. 1f**). Since CD25 is also a marker of CD4+ regulatory T (T_reg_) cells^35^, we also measured the proportion of CD4+ CD25+ FoxP3+ T_reg_ cells in each condition. T_reg_ cells comprised <5% of the expanded T cells in all conditions (**Supplementary Fig. 1e**), indicating that the conditions used in this study do not support the proliferation of FoxP3+ T_reg_. We also evaluated the expression of PD-1 as an indicator of potential T cell exhaustion since its expression on expanded CAR T cells is correlated with poorer patient outcomes^36^. PD-1 expression was generally low on cells from all conditions except for CD4+ REP cells where 50% of the CD4+ T cell population expressed PD-1 (**Fig. 1g**). Moreover, upon reactivation, REP expanded T cells displayed higher TNFα and IFNγ production (**Fig. 1h**). Collectively, these results establish that T cell products vary in proliferative capacity, differentiation (T_EFF_ and T_EM_), function (TNFα and IFNγ) and expression of exhaustion makers (PD-1).

### Mitochondria mass, activity and ROS vary across all conditions

Mitochondrial activity and biomass have been recognized in both pre-clinical and clinical models to be indicators of improved CAR T cell therapeutic efficacy. Specifically, low mitochondrial activity and increased mitochondrial biogenesis supports T cell persistence and anti-tumor function in chronic lymphocyte leukemia patients^37^. Therefore, we assessed the impact of expansion conditions on mitochondrial activity and mitochondrial mass as determined by flow cytometry analysis of cells stained with MitoTracker Deep Red and MitoTracker Green, respectively. Overall, there was no consistent pattern that emerged across conditions although statistical differences were observed depending on the pair-wise comparisons (**Fig. 2a,b and Supplementary Fig. 2a**). We found that the REP expanded T cells produced the highest cell number also had the highest mitochondrial activity (**Fig. 2a**). Notably, high mitochondrial activity is associated with a terminal differentiation state and poor tumor killing capacity^38^. In contrast, T cells maintained in ICM had the second largest expansion with lower mitochondrial activity compared to the REP, but had significantly higher mitochondrial mass compared to all other conditions (**Fig. 2b**). The substantial increase in mitochondrial biogenesis observed in the ICM condition was not due to a difference in cell size (**Supplementary Fig. 2b**).

**Fig. 2:**
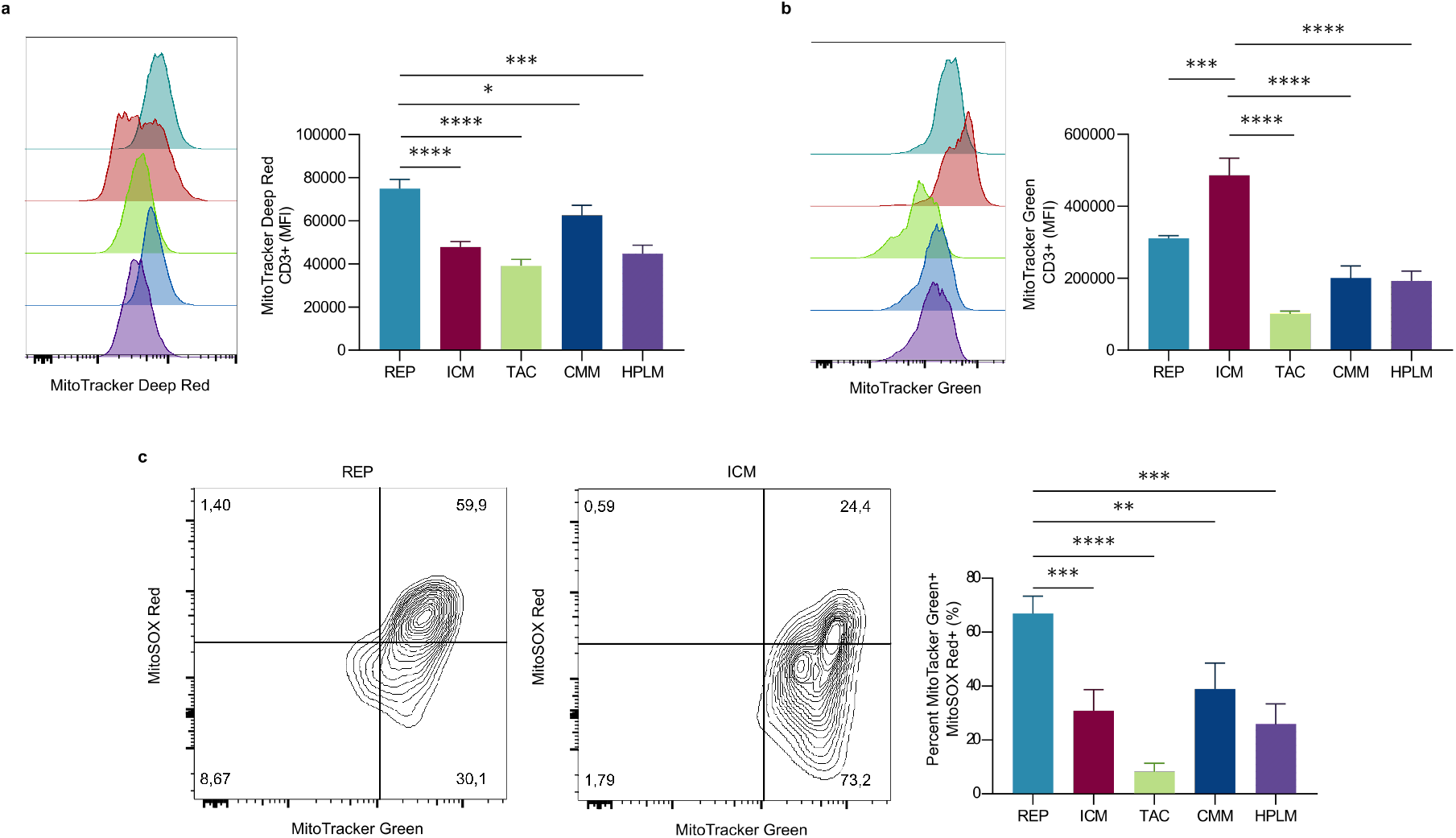
T cell products exhibit different mitochondrial phenotypes. **(a-c)** T cells from three healthy donors were expanded in 5 different conditions for 12 days. **(a,b)** Representative plot (left) and tabulated data (right) for median fluorescence intensity (MFI) of mitochondrial activity (MitoTracker Deep Red) and **(b)** mitochondrial mass (MitoTracker Green) **(c)**. Representative plot (left) and tabulated data (right) for percentage of mitochondrial mass high and mitochondrial ROS high (MitoSOX) live populations. Bar graphs represent mean of n=3 +SEM from healthy donors. Statistical significance was calculated by one-way ANOVA (* p <0.05, ** p <0.01, *** p <0.001, **** p <0.0001).

Cells with elevated levels of mitochondrial activity often results in elevated mitochondrial reactive oxygen species (ROS) production that can limit the cells functional capability in conditions of oxidative stress such as the TME. Therefore, we investigated how the mitochondrial activity is associated with mitochondrial ROS for each expansion product. We found that mitochondrial activity and mitochondrial ROS were positively correlated in all conditions, although at varying degrees (**Supplementary Fig. 2c**). In accordance with the elevated mitochondrial activity, REP expanded T cells had the highest ROS production, where approximately 70% of mitochondria were positive for ROS (**Fig. 2c**). In contrast to the REP, the ICM condition displayed higher mitochondrial mass and low levels of ROS (**Fig. 2c**). While the conditions with the lowest proliferation and mitochondrial activity, TAC and HPLM appeared to produce the lowest mitochondrial specific ROS.

### Expansion conditions dictate nutrient uptake but is uncoupled from T cell function

Media conditions are often proprietary, so the precise composition of basic nutrient levels are not available. Therefore, the concentration of two key nutrients, glucose and glutamine, was assessed as both are well studied in terms of their regulation of T cells^39–41^. The concentrations of glucose and glutamine varied significantly across conditions, 5.92-27.25 mM and 0.76-6.68 mM, respectively (**Table 1**). Thus, we investigated how the variation in nutrient levels influenced uptake between days 11-12. All five conditions supported varying levels of both glucose and glutamine uptake (**Fig. 3a,b and Supplementary Fig. 3a,b**). Overall, conditions with elevated glucose concentrations tended to have increased uptake (**Supplementary Fig. 3a**), while glutamine concentrations did not appear to be associated with the level of uptake (**Supplementary Fig. 3b**). Furthermore, glucose uptake appeared to be independent of proliferation as both the highest (REP, ICM CMM) and lowest (HPLM) proliferative conditions had significant changes in glucose concentration compared to their respective fresh media condition (**Fig. 3a**). The elevated nutrient uptake observed in HPLM expanded cells may be due to the differing proliferative state at day 11, as they were most proliferative at day 11 **(Supplementary Fig. 3c)**. Furthermore, despite containing supraphysiological level of glucose and glutamine, the REP expanded T cells resulted in no significant change in glutamine concentration in the media over 24 hours of culture, indicative of low glutamine uptake and a preference for glucose as a carbon source. In contrast, the CMM expansion condition produced significant differences in both glucose and glutamine concentrations in the media after 24 hours (**Fig. 3a,b**). To gain insight into whether the dependence on glucose in the REP was mediated by the cell culture media formulation, we expanded T cells using the respective 4 conditions and on day 11 replaced the media with the REP media (CTL:AIM-V) for 24 hours. This switch resulted in a significant change in glucose concentration in CTL:AIM-V with a concomitant no change in glutamine across almost all expansion products (**Fig. 3c,d**). Thus, this media replacement phenocopied the glucose metabolism of the REP expanded condition in the CTL:AIM-V. Although there was a modest reduction in glutamine consumption by CMM expanded T cells that had been switched to CTL:AIM-V, the shift to reduced glutamine uptake was the most dramatic of all conditions. These findings imply that T cells exhibit metabolic flexibility and adapt to their surrounding nutrient environments, regardless of their original metabolic state at the time of activation and/or expansion.

**Fig. 3:**
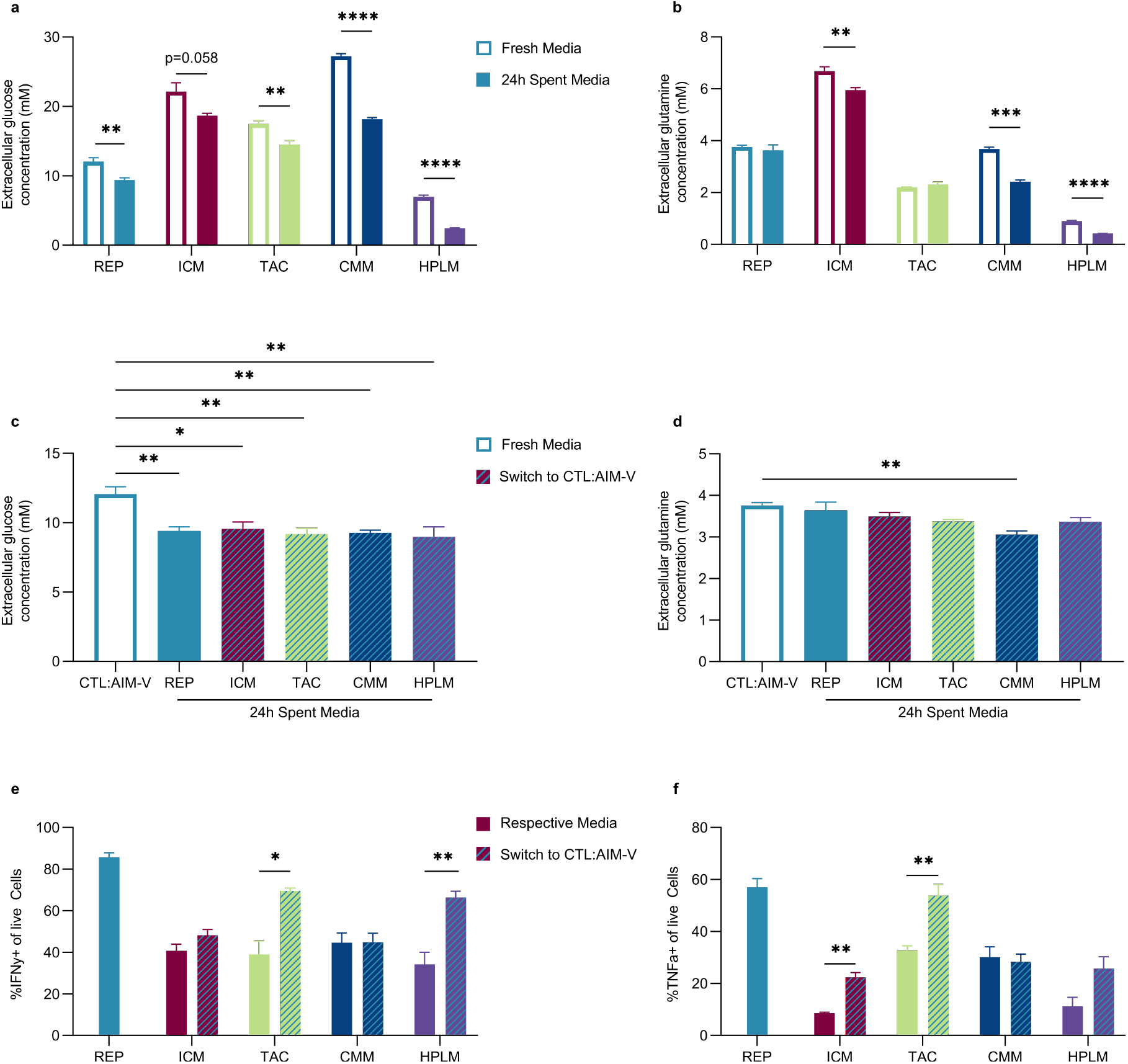
Expansion conditions dictate nutrient uptake but is uncoupled to T cell function. **(a-d)** T cells from three healthy donors were expanded in 5 different conditions for 11 days. Glucose and glutamine concentrations in the media were measured between days 11-12. **(a)** Extracellular glucose and **(b)** glutamine concentrations in the fresh media (white bars) and spent media after culture for 24 hours (solid bars). **(c, d)** On day 11, expanded T cells from all 5 conditions were switched to the REP media (CTL:AIM-V) for 24 hours. **(c)** Extracellular glucose and **(d)** glutamine concentrations in fresh CTL:AIM-V (white bar), and spent media after culture for 24 hours with REP expanded cells (blue bar) and the four other expansion products in CTL:AIM-V (blue dashed bar). **(e,f)** On day 12, T cells from all 5 conditions underwent CD3/CD28 reactivation for 2 days in their respective conditions (solid bars) or were switched to CTL:AIM-V (blue dashed bars). Percentage of **(e)** IFNγ and **(f)** TNFα positive cells. Bar graphs represent mean n=3 +SEM from healthy donors. Statistical significance was calculated by one-way ANOVA (**a,b,e,f**) or Student’s t-test (**c,d**) (* p<0.05, ** p<0.01, *** p<0.001, **** p<0.0001).

To uncover if the shift in metabolism in CTL:AIM-V was also linked to increased function as observed in the REP expanded T cells, we reactivated all products in their respective conditions or in CTL:AIM-V for 2 days. Unlike glucose and glutamine uptake, T cell function was not universally influenced by the change to CTL:AIM-V media across conditions (**Fig. 3e,f**). Although all T cell products were reactivated in CTL:AIM-V, differences in IFNγ and TNFα production was observed. While TAC significantly increased in both IFNγ and TNFα production in CTL:AIM-V, CMM cells displayed no change in either cytokine (**Fig. 3e,f**). This demonstrates that while metabolism adapts to the change in extracellular conditions, the inherited impacts of the initial expansion conditions on T cell function cannot be universally overcome by switching conditions.

### Glucose utilization and phenotype is influenced by media

To gain further insight into the influence of media on T cell metabolism, we investigated the difference in glucose utilization pathways between the REP and ICM condition. These conditions produced the largest cell expansions with similar glucose uptake levels regardless of glucose concentrations in the media (**Table 1 and Supplementary Fig. 4a**). Therefore, stable isotope labeling was performed using [U-^13^C]glucose to delineate how carbon utilization differed in REP and ICM conditions (**Fig. 4a**). Current *in vitro* models suggest that the rate of glycolysis is matched to the rate of proliferation. Indeed, we found that aerobic glycolysis was active in both conditions, producing high extracellular and intracellular lactate M+3 fractions in both CD4+ and CD8+ T cells (**Fig. 4b and Supplementary Fig. 4b)**. However, the REP condition produced significantly more extracellular lactate compared to ICM (**Supplementary Fig. 4c**), consistent with a more glycolytic and activated phenotype (**Fig. 1f,g**). Furthermore, the ICM condition produced roughly 30% less extracellular glucose derived lactate M+3, indicating that other carbon sources support lactate production (**Fig. 4b**). To assess if glucose utilization was dependent on the culture medium, the day 11 ICM expanded T cell products were placed in CTL:AIM-V. After 24 hours in CTL:AIM-V, the ICM expanded T cell products did not increase in lactate production (**Supplementary Fig. 4c**), however their carbon utilization pathways shifted resulting in a significantly higher fraction of lactate M+3 similar to the REP condition (**Fig. 4b**). A similar enrichment in the ^13^C labeling pattern was observed for alanine M+3. There was a higher level of alanine M+3 enrichment in the REP and ICM switched to CTL:AIM-V compared to ICM cells in their respective condition (**Fig. 4c**). Interestingly, ICM contains approximately 7 times more alanine than CTL:AIM-V (**Supplementary Fig. 4d**), which was associated with higher intracellular levels (**Supplementary Fig. 4e**). Therefore, the reduced fraction of alanine M+3 in the ICM condition could be due to a combination of increased *de novo* production and/or uptake.

**Fig. 4:**
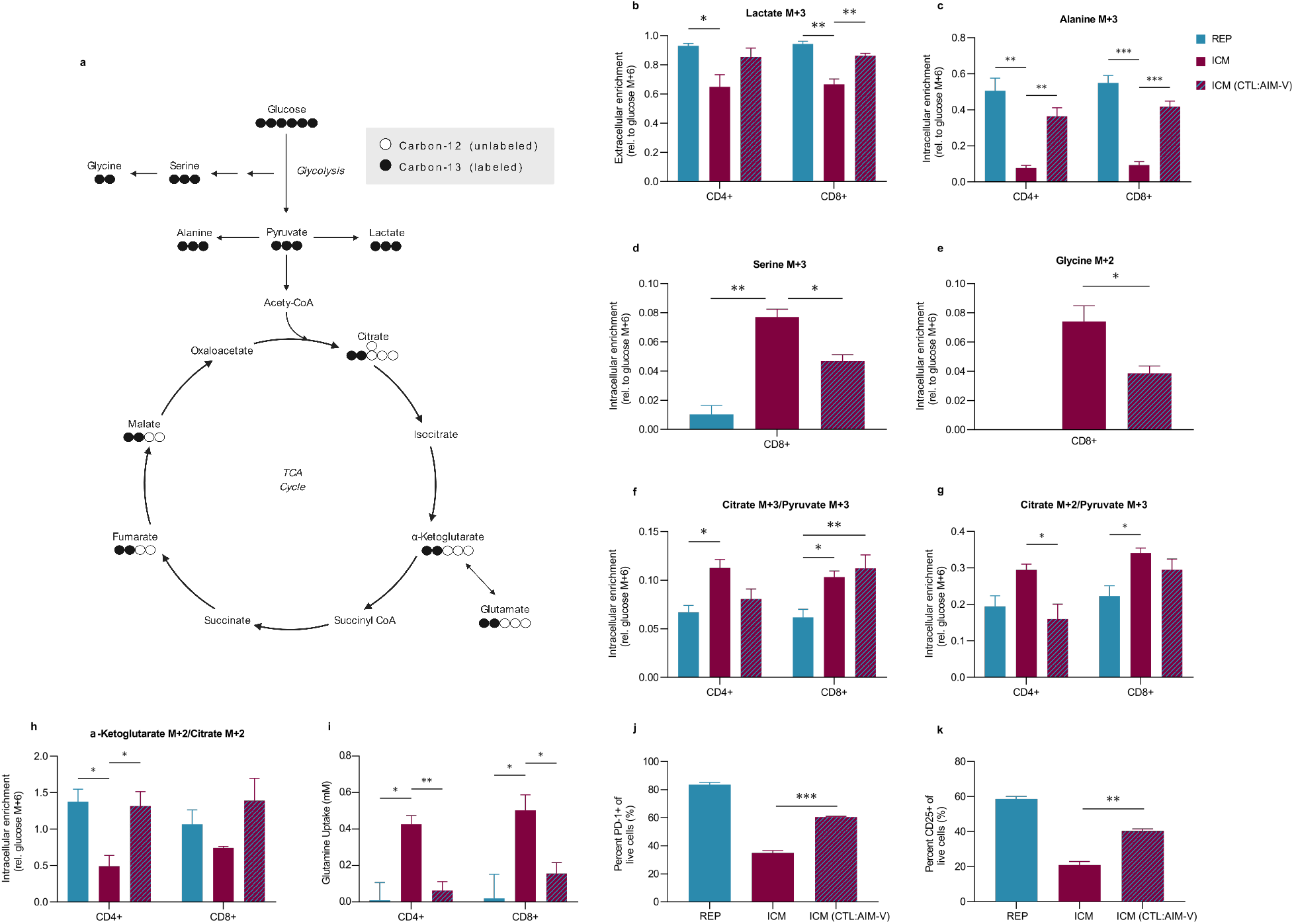
Culture media influences glucose utilization and the metabolic profile of T cells. **(a)** Schematic of [U-^13^C]glucose metabolism, circles represent the carbons for each metabolite. **(b-k)** Three healthy donors were expanded in REP and ICM for 11 days. On day 11, CD4+ and CD8+ cells were isolated and incubated in the following [U-^13^C]glucose conditions for 24 hours: REP cells in CTL:AIM-V (blue bar), ICM cells in ImmunoCult-XF (red bar) and ICM cells cultured in CTL:AIM-V (red and blue dashes). **(b)** Extracellular lactate M+3 relative to extracellular glucose M+6 enrichment. **(c)** Intracellular alanine M+3 relative to intracellular glucose M+6 enrichment. CD8+ intracellular enrichment of **(d)** serine M+3, and (**e)** glycine M+2 relative to intracellular glucose M+6 enrichment. **(f)** Pyruvate carboxylase activity (Citrate M+3/Pyruvate M+3) and **(g)** Pyruvate dehydrogenase activity (Citrate M+2/Pyruvate M+3). **(h)** Proportion of intracellular α-ketoglutarate M+2 to citrate M+2 enrichment. **(i)** Glutamine uptake (mM) from day 11-12 in CD4+ and CD8+ T cells. Percentage of PD-1 **(j)** and CD25 **(k)** positive cells. Data are shown as mean of n=3 +SEM from healthy donors. Statistical significance was calculated by Student’s t-tests (* p<0.05, ** p<0.01, *** p<0.001).

Due to the decrease in lactate and alanine labeling in ICM conditions, we interrogated other upstream glucose-derived metabolites. Serine is a non-essential amino acid that contributes to nucleotide biosynthesis. Recent evidence indicates that serine is necessary to support CD8+ T cell expansion and effector function^42,43^. The [U-^13^C]glucose metabolite isotope analysis revealed a greater fractional enrichment of labeled serine and glycine in ICM expanded T cells compared to REP expanded T cells. More specifically, CD8+ T cells from the ICM expansion had significant enrichment of serine M+1 and M+3 and glycine M+2 compared to CD8+ T cells from the REP expansion (**Fig. 4d,e and Supplementary Fig. 4f**). Although the relative serine labeling was reduced in ICM expanded T cells that were switched to the CTL:AIM-V, there were detectable levels of serine M+3 enrichment. This suggests that relative to the REP expansion conditions, ICM expansion conditions promote one-carbon metabolism pathways in CD8+ T cells and that changes in one-carbon metabolism may be less susceptible to fluctuations in the levels of extracellular nutrients.

At the peak of an effector response, T cells undergo a metabolic switch from glycolysis to OXPHOS supporting mitochondrial biogenesis and T cell memory development. Therefore, we assessed the fate of the [U-^13^C]glucose carbons into the mitochondria (**Fig. 4a**). Citrate M+2/Pyruvate M+3 and Citrate M+3/Pyruvate M+3 ratios serve as surrogates for pyruvate dehydrogenase (PDH) and pyruvate carboxylase (PC) activity respectively^44,45^. Between conditions, both CD4+ and CD8+ T cells from the ICM expansion had higher PC and PDH activity than CD4+ and CD8+ T cells from the REP expansion (**Fig. 4f,g**). The ICM-mediated contribution of glucose-derived carbons into the TCA cycle is consistent with the observed increase in mitochondrial biogenesis (**Fig. 2b**). However, CD4+ T cells seem to be more influenced by the switch to CTL:AIM-V condition than the CD8+ T cells which maintained the PDH and PC labeling patterns regardless of condition. Furthermore, the ICM expanded T cells diverted roughly 50% of α-ketoglutarate M+2 from citrate M+2 (**Fig. 4h**). This reduction in α-ketoglutarate M+2 was corroborated by the reduction in glutamate M+2 and malate M+2 fractional enrichment in the ICM expansion condition (**Supplementary Fig. 4g,h**). The loss in α-ketoglutarate M+2 enrichment was likely due to entry of unlabeled carbons from glutamine catabolism, both of which were reversed in CTL:AIM-V conditions (**Fig. 4h,i**). These results imply that REP and ICM expansion conditions differentially control the source of carbons entering the TCA cycle. ICM expanded T cells direct glucose-derived carbons preferentially into citrate via PC and PDH, and likely use glutaminolysis as an alternative carbon source to support synthesis of TCA cycle metabolites.

The REP condition supports increased glycolysis, glucose dependence, elevated mitochondrial activity and ROS generation, many of which are hallmarks of T cell exhaustion. Indeed the REP cells resulted in greater than 50% of the expanded T cells expressing PD-1 (**Fig. 1g**). Therefore, we investigated if conditioning ICM expanded T cells in CTL:AIM-V would also influence exhaustion. On day 12, REP and ICM expanded T cells were reactivated in their respective conditions or in CTL:AIM-V for an additional 2 days. The REP expanded T cells showed a significantly higher proportion of cells that were PD-1 and CD25 positive compared to ICM expanded T cells (**Fig. 4j,k**). However, the ICM expanded cells that were reactivated in CTL:AIM-V conditions also significantly increased in activation and exhaustion markers resembling the exhaustion state of the REP expanded T cells (**Fig. 4j,k**). This suggests that while ICM expanded T cells do not increase glycolysis when switched to CTL:AIM-V, the nutrient conditions in the CTL:AIM-V media resulted in changes to both glucose and glutamine metabolism that were sufficient to impart alterations in activation and exhaustion phenotypes.

### The tumor microenvironment imposes different metabolic constraints on T cell products

Given the observed changes in T cell metabolism and phenotypes when expanded T cells are placed into different media, it is possible that T cells manufactured under these parameters may also be affected by the TME. There is a growing appreciation that *ex vivo* clinical manufacturing conditions used to expand T cells for adoptive cell therapies may impact the effectiveness of the final T cell product when the infused T cells encounter the nutrient depleted TME^46,47^. To test this possibility, patient-derived ovarian cancer ascites fluid was used as a proxy for the TME. We expanded T cells in all five conditions and on day 12 of the expansion, T cell products were cultured and reactivated in 100% ascites fluid for 2 days (**Fig. 5a**).

**Fig. 5:**
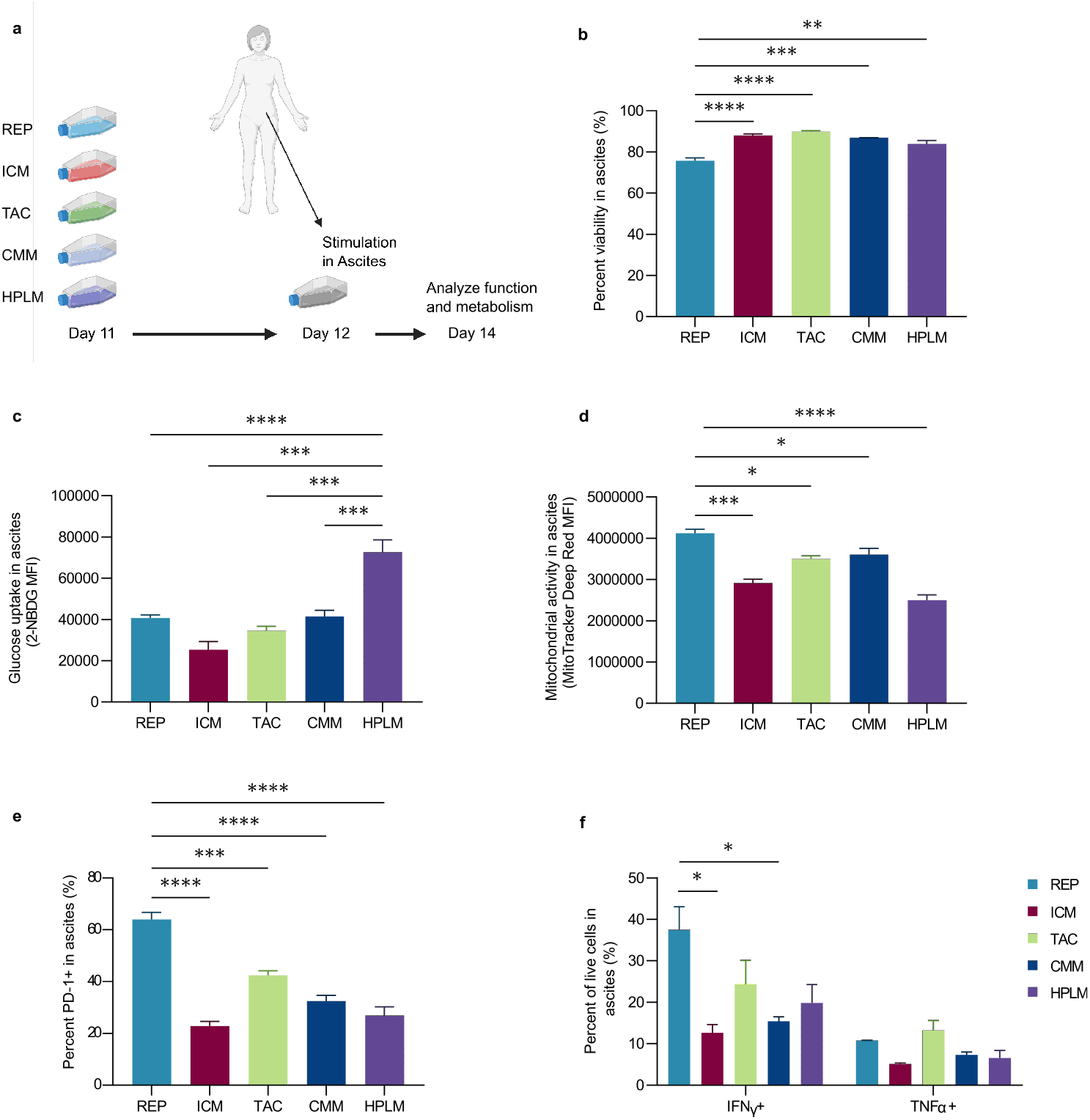
T cell products exhibit different metabolic and functional phenotypes in the ascites tumour microenvironment. **(a-f)** T cells from three healthy donors were expanded in 5 different conditions over 12 days. On day 12 T cell products were reactivated (CD3/CD28) in ascites, T cell metabolism and function was assessed after 2 days. **(a)** Schematic of experimental timeline. **(b)** Percentage of live cells in ascites supernatant. Median fluorescence intensity (MFI) of **(c)** glucose uptake (2-NBDG) and **(d)** mitochondrial activity (Mitotracker DeepRed) in the ascites. **(e)** Percentage of PD-1, **(f)** IFNγ and TNFα positive cells in the ascites. Data are shown as mean of n=3 +SEM from healthy donors. Statistical significance was calculated by one-way ANOVA (* p<0.05, ** p<0.01, *** p<0.001, **** p<0.0001).

Four of the five expanded T cell conditions maintained their viability in the presence of ascites (**Fig. 5b**). However, the REP expanded T cells showed a consistent and noticeable decrease in viability compared to the remaining four conditions.

The ascites environment has been reported to be immune suppressive owing to reduced levels of critical nutrients such as glucose and glutamine^48,49^. Consistent with this, analysis of the ascites supernatant revealed significantly lower levels of both glucose and glutamine compared to levels in all of the media formulations except for HPLM (**Table 1 and Supplementary Fig. 5a,b**). The HPLM was developed to more closely resemble the metabolic conditions in normal human plasma. Using the fluorescent glucose analog 2-NBDG, we found that despite containing similar levels of glucose to that found in the ascites, the HPLM expanded T cells had significantly higher 2-NBDG uptake in ascites (**Fig. 5c**). The other four conditions showed no change in glucose uptake between their respective cultured conditions and ascites fluid (**Supplementary Fig. 5c**). In contrast to the lack of changes in glucose uptake, mitochondrial metabolism was lower in all of the expanded T cell products in ascites fluid (**Supplementary Fig. 5d**).

However, similar to what was seen in the media conditions, the REP had the highest mitochondrial activity in ascites fluid compared to the other conditions (**Fig. 5d**). Although the REP condition had high levels of PD-1 expression in the ascites across all conditions (**Fig. 5e**), the expression of PD-1 was reduced in the ascites (**Supplementary Fig. 5e**). Since PD-1 is also a marker of activation, reduced PD-1 expression could be a reflection of the suppressed activation state of T cells in ascites. Indeed, the T cell activation marker CD25 was also reduced across conditions in the ascites further supporting the contention that ascites is an immune suppressive environment (**Supplementary Fig. 5f**). Consistent with the reduction in mitochondrial activity, expanded T cells cultured in ascites resulted in a decrease in IFNγ and TNFα production regardless of the expansion condition (**Supplementary Fig. 5g,h**). While the REP expanded T cells produced the largest proportion of IFNγ positive cells, there was no significant difference in TNFα expression across conditions in ascites fluid (**Fig. 5f**).

## DISCUSSION

Clinical grade T cells are manufactured under stringent and defined conditions yet to date there is no universal media formulation or standardized protocols in the field. Moreover, the release criteria do not take into account how different expansion conditions alter T cell metabolism or how these conditions may impact T cell function post-infusion. Recent work has highlighted important differences in how T cells metabolize glucose *in vitro* versus *in vivo*^20^. During CD8+ T cell responses to physiological infection, T cells adopt an oxidative metabolic phenotype with greater flux of glucose-derived carbons into serine and other biosynthetic pathways. This is consistent with another study showing that glucose pre-conditioning can metabolically enhance adoptive T cell anti-tumor immunity potentially by shifting cells into a temporary oxidative state^47^. These studies suggest that the nutrient conditions during the *ex vivo* manufacturing could be tailored to enhance the fitness of T cells following infusion into patients.

We evaluated how five commonly used *ex vivo* expansion culturing conditions alter the metabolic state and phenotypes of CD4+ and CD8+ T cells. Across these conditions, there were pervasive differences in metabolism that resulted in shifts in differentiation and function. We used the REP formulation as a benchmark given its early use in the first adoptive T cell trials for the treatment of late-stage metastatic melanoma. Although all conditions resulted in T cell proliferation, the extent of expansion was markedly different and ranged from 40 to 860-fold. However, the baseline concentration of glucose present in each media did not correlate with the degree of expansion. The CTL:AIM-V media used in the REP contained 11 mM glucose but yielded the highest fold expansion despite the ICM, TAC and CMM media containing twice the concentration of glucose. Although REP and ICM T cells resulted in a similar level of expansion, [U-^13^C]glucose isotope tracing studies revealed that T cells expanded in the ICM culture medium had lower levels of extracellular lactate and lactate M+3 enrichment, suggesting that glycolysis was less engaged in these cells. The reduction in lactate was associated with a greater proportion of CD8+ T_EFF_ cells and decreased the proportion of T_EM_. This observation was unexpected given that T_EFF_ cells have been shown to be highly glycolytic and the transition to T_EM_ cells is associated with a switch to oxidative metabolism. These results suggest that during *ex vivo* expansion, metabolites other than glucose that are found in the ICM media play a role supporting T_EFF_ cell differentiation and proliferation. Moreover, the differences in proliferation as well as differentiation phenotypes could not be fully attributed to the metabolism of T cells implying that these phenomena are disconnected under the *ex vivo* conditions tested in our study. It is also possible that the specific TCR stimulation method activates unique signaling programs that could contribute to the different metabolic responses. However this was not observed in the CMM and HPLM conditions. Furthermore, this is independent from the TCR clonality as no major biases were seen in the outgrowth of TCR diversity.

Due to the implications of mitochondrial metabolism in the context of CAR and TIL therapy, we assessed the mitochondrial phenotype produced from all 5 conditions. ICM expanded T cells had significantly higher mitochondrial mass compared to all other conditions. This was surprising as the majority of ICM cells had a T_EFF_ phenotype and increases in mitochondrial biogenesis is commonly associated with T memory subsets. The REP expanded T cells also showed high mitochondrial activity, which correlated with increased production of mitochondrial specific ROS. Although ROS production is required for T cell activation, elevated production of free radicals can induce mitochondrial stress and trigger cell apoptosis. This mitochondrial phenotype may explain the poor persistence *in vivo*, and poor survival and cytotoxic efficacy under oxidative stress, an observation that has been previously reported for T cells expanded using the REP formulation^50^. In contrast, the ICM medium produced cells with increased mitochondrial mass and reduced activity and ROS production compared to the REP. Based on previous clinical findings, a product with this mitochondrial profile may be most advantageous for cell based therapies^37^. However, in our TME model used here, no expansion condition supported sustained mitochondrial activity in the presence of ascites fluid.

To further understand how culture medium influences T cell metabolism, we traced [U-^13^C]glucose utilization in T cells that were expanded using the REP and ICM condition, as well as ICM expanded T cells and subsequently switched into CTL:AIM-V. These results revealed that the elevated glycolysis labeling and reduced of TCA cycle intermediate labeling was media dependent. Of note, ^13^C enrichment was reduced in metabolites downstream of citrate M+2 in ICM expanded T cells compared to REP T cells and ICM T cells switched in CTL:AIM-V for 24 hours. This is likely due to the increased uptake of glutamine in ICM conditions, while REP and T cells switched to CTL:AIM-V media preferentially utilize glucose over glutamine as a carbon source. These patterns were also associated with a corresponding increase in both intracellular and extracellular alanine M+3. Alanine has been shown to be essential for T cell activation and can be produced from a transamination reaction with pyruvate and glutamate via alanine aminotransferase. However, this increased proportion of fractional enrichment of alanine M+3 is also a reflection of the lower proportions of unlabeled extracellular and intracellular alanine in T cells expanded under CTL:AIM-V compared to T cells expanded under ICM media. When ICM expanded T cells were switched into CTL:AIM-V, the total intracellular alanine levels decrease (**Supplementary Fig. 4e**) supporting the idea that REP expanded T cells may have reduced alanine uptake. This implies that exogenous alanine in the ICM medium is sufficient to meet cellular demands, while T cells expanded with the REP medium appear to require synthesis of alanine. However, further studies are needed to formally demonstrate this possible scenario.

Isotope tracing studies also revealed metabolic pathways that are less susceptible to reprogramming due to fluctuations in extracellular nutrient conditions. For example, some metabolic phenotypes of the ICM expanded T cells persisted even when the T cells were switched to CTL:AIM-V medium, including the increased PDH and PC activity in CD8+ T cells and enrichment of ^13^C to one carbon metabolism pathways. These data provide an example of how T cells adapt their metabolism based on extracellular levels of nutrients and the plasticity of certain metabolic pathways. This also further supports the idea that some metabolic pathways may have a more permanent inheritability throughout the expansion, while the utilization of other metabolic pathways such as glycolysis may change rapidly depending on precise conditions at the time. It will be important to consider which metabolic pathways and cell phenotypes will be maintained after the expanded T cells are administered and subsequently traffic into the TME.

Lastly, we tested how all five expansion conditions supported T cell metabolism and function when transferred to patient-derived ovarian cancer ascites fluid, a known immunosuppressive TME. Due to the elevated level of ROS and PD-1 expression in the REP expanded T cells, we speculated that this state may contribute to activation-induced T cell death upon reactivation in ascites^52^. As expected, the REP condition had significantly reduced T cell viability after reactivation compared to the other media conditions. Interestingly, glucose uptake was not suppressed across all conditions and actually increased in T cells expanded under the HPLM conditions. This result further highlights the notion that the *ex vivo* metabolism of T cells is not necessarily coupled to their functional behavior when subjected to the TME. The increase in glucose uptake could be associated with higher CD28 expression as it has been shown that CD28 supports Akt activation^53^. It is tempting to speculate that HPLM may be more suited for expansion of second or third generation CAR T cells that encode for CD28 co-stimulatory domains versus 4-1BB co-stimulatory domains. HPLM contains similar glucose levels to that found in ascites suggesting that initial expansion of T cells in low glucose conditions programmed T cells to subsist under environments where glucose levels are limited^47^. On the other hand, mitochondrial activity was significantly suppressed across all conditions, which is commonly observed in the TME and could contribute to the observed suppression of T cell function. Similar to other reports, we observed that T cell products with elevated mitochondrial activity showed a tendency to have higher PD-1 expression^52^. This demonstrates that the impact of the media formulations extends beyond nutrient uptake and glucose utilization in that they directly affect T cell activation.

In conclusion, we uncovered distinct metabolic programs activated by *ex vivo* clinical expansion protocols. The observed differences under various media formulations contributed to the skewed patterns of T cell differentiation and effector function, outputs that could be uncoupled with their metabolic profiles. Most T cell-based immunotherapies focus on applications to improve cell expansion and to produce phenotypes that enable *in vivo* persistence and maximal cytolytic function. Here we demonstrate that media formulations may influence metabolic fidelity of the final immunotherapeutic product. Further studies will be required to determine whether these metabolic states imparted by different *ex vivo* expansion conditions ultimately impact the *in vivo* anti-tumor ability of T cells and their long-term persistence.

## METHODS

### Cell culture reagents

Five media conditions were interrogated in this study. (1) Complete CTL:AIM-V media, used in rapid expansion T cell protocol (REP), consisted of equal parts of supplemented RPMI 1640 and AIM-V media. To a 1X RPMI 1640 Medium basal, the following components were added: 2 mM L-Glutamine (Fisher), 10% heat-inactivated (56°C, 60 min) human AB serum (Sigma), 12.5 mM HEPES (Fisher), 1X Penicillin Streptomycin solution (Fisher), and 50 μM β-Mercaptoethanol (Sigma). AIM-V Medium (Invitrogen) was supplemented with 20 mM HEPES and 2 mM L-glutamine (CTL:AIM-V). (2) ImmunoCult™-XF T Cell Expansion Medium (STEMCELL Technologies) was supplemented with 1X Penicillin Streptomycin solution (ICM). (3) TexMACS™ medium (Miltenyi) was supplemented with 3% heat-inactivated human AB serum (Sigma) and 1X Penicillin Streptomycin solution (TAC). (4) Corning media consisted of Lymphocyte Serum-Free Medium KBM 581 (Corning) supplemented with 3% heat-inactivated human AB serum (CMM). (5) Basal Human Plasma-like Medium (HPLM) (kindly provided by Dr. Jason Cantor)^21^ was prepared with four additional components (5 μM acetylcarnitine, 5 μM α-ketoglutarate, 5 μM malate, and 3 μM uridine) as recently reported^23^, and then further supplemented with 3% heat-inactivated human AB serum (HPLM). All complete media were filtered through a 0.22 μM filter prior to use.

### Patient ascites collection

Patient specimens and clinical data were obtained through the BC Cancer Tumour Tissue Repository, certified by the Canadian Tissue Repository Network. All specimens and clinical data were obtained with either informed written consent or a formal waiver of consent under protocols approved by the Research Ethics Board of BC Cancer and the University of British Columbia (H07-00463). Patient ascites were centrifuged at 1500 rpm for 10 min at 4°C to pellet cells and supernatant was frozen at -80°C. The preserved supernatants were thawed for the functional assays as described below.

### T cell expansions

PBMCs were isolated from six human peripheral blood leukapheresis packs (STEMCELL Technologies) using Ficoll gradient density centrifugation. CD3+ T cells were isolated from cryopreserved PBMCs using human CD3 MicroBeads (Miltenyi) according to the manufacturer’s instructions. Specific details about stimulation protocols used can be found in **Table 1**. All media were supplemented with 300 IU/ml IL-2 (Novartis) before cell culture. REP CD3+ T cells (1.0 × 10^5^) were stimulated with 50 Gy-irradiated feeder PBMCs (2.0 × 10^7^) and soluble CD3 (30 ng/ml; OKT3) in complete CTL:AIM-V media. Other expansion methods were seeded using 1.0 × 10^6^ CD3+ T cells in 1 ml of media in a 48-well plate. ICM T cells were stimulated with ImmunoCult™ Human CD3/CD28 T Cell Activators (25 μl/ml, STEMCELL Technologies). TAC T cells were stimulated with T Cell CD3/CD28 TransAct™, human (10 μl/ml, Miltenyi). CMM and HPLM T cells were stimulated with plate-bound CD3 (5 μg/ml; OKT3) and soluble CD28 (2 μg/ml; 15EB). On day 2-3, the media from TAC cells was replaced with fresh media as per manufacturer’s instructions. T cells were first split either on day 3 (ICM) or day 4 (other conditions) of the expansion, and subsequently split as needed to maintain a concentration of 100,000-600,000 cells/ml (ICM) or 500,000-1,000,000 cells/ml (other conditions). During the expansion, cells were isolated, diluted in trypan blue, and counted using a hemocytometer. Cell counts were measured throughout the expansion to ensure protocol-recommended densities were maintained. T cell reactivation in their respective condition, or ascites, was performed using plate-bound CD3 (5 μg/ml; OKT3) and soluble CD28 (2 μg/ml; 15EB) for 2 days.

### Phenotypic and metabolic profiling by flow cytometry

Prior to expansion (day 0), cells were collected from CD3+ magnetic bead isolation. Cells were stained with viability dye (eFlour506, Thermo) diluted in PBS (Invitrogen) at 4°C for 15 minutes. Cells were stained with a panel of antibodies (**Supplementary Table 1**) in flow cytometry staining buffer for 30 minutes at 4°C. After staining, cells were washed twice and resuspended in flow cytometry staining buffer prior to flow cytometry analysis. Following expansion, separate flow cytometry panels were used on day 12, for cell phenotyping and mitochondrial analysis (**Supplementary Table 1**). For mitochondrial analysis, cells were stained with MitoTracker Deep Red (10 nM), MitoTracker Green (100 nM) or MitoSOX Red (2.5 uM) for 30 minutes at 37°C. Cells were then washed twice with PBS, and then stained with viability dye at 4°C for 15 minutes. For cell phenotype analysis, cells were then washed with flow cytometry staining buffer before being fixed and permeabilized using Fixation/Permeabilization Solution Kit (Biosciences) as per the manufacturer’s instructions. Cells were stained with a panel of antibodies (**Supplementary Table 1**) in 1X BD Perm/Wash™ buffer (Biosciences) for 30 minutes at 4°C. After staining, cells were washed twice in 1X BD Perm/Wash™ buffer (Biosciences) and resuspended in flow cytometry staining buffer prior to flow cytometry analysis. Flow cytometry analysis was carried out using a Cytek Aurora spectral flow cytometer (3L-16V-14B-8R configuration). Data were unmixed using SpectroFlo Software (Cytek), and manually gated and analyzed using FlowJo V10.6.1. Figures were created using GraphPad Prism 8.1.2. To assess phenotype, cells were classified as naïve (T_N_; CCR7+CD45RO–), effector (T_EFF_; CCR7– CD45RO–), effector memory (T_EM_; CCR7–CD45RO+), or central memory (T_CM_; CCR7+CD45RO+)^24^. To assess effector function following expansion, on day 12 in their respective condition or ascites, T cells were reactivated for 2 days using plate-bound CD3 (5 μg/ml; OKT3) and soluble CD28 (2 μg/ml; 15EB). Six hours prior to staining, cells were treated with BD GolgiStop™ (1 μl/ml, Biosciences) to assess TNFα and INFγ production. Cells that underwent metabolic analysis (**Supplementary Table 1**) with 2-NBDG (100 uM) and MitoTracker Deep Red (10 nM) were not treated with BD GolgiStop™.

### Vβ spectratyping by flow cytometry

To assess T cell receptor (TCR) diversity following expansion (day 12), cell surface TCR Vβ repertoires were profiled using the IOTest Beta Mark Kit (Beckman Coulter) as per the manufacturer’s guidelines. Cells were washed in PBS (Invitrogen) and blocked with Anti-Hu Fc Receptor Binding Inhibitor (eBiosciences) for 10 minutes at room temperature. Cells were stained with a panel of antibodies (**Supplementary Table 1**) in flow cytometry staining buffer with BD Horizon Brilliant Stain Buffer Plus (BD Biosciences) for 30 minutes at 4°C. Cell viability was assessed using the Zombie NIR™ Fixable Viability Kit (Biolegend) as per the manufacturer’s guidelines. After staining, cells were washed once in flow cytometry staining buffer and resuspended in flow cytometry staining buffer prior to flow cytometry analysis using a Cytek Aurora spectral flow cytometer (3L-16V-14B-8R configuration). Data was unmixed using SpectroFlo Software (Cytek), manually gated and analyzed using FlowJo V10.6.1, and figures were created using GraphPad Prism 8.1.2. Manual gating was carried out following the manufacturer’s guidelines (Beckman Coulter).

### Isotope tracing analysis by GC-MS using [U-^13^C]glucose

For metabolic labeling experiments, complete CTL:AIM-V labeling medium consisted of glucose-free CTL:AIM-V with 5% heat-inactivated human AB serum supplemented with 11 mM uniformly labeled ^13^C-glucose ([U-^13^C]glucose) (Cambridge Isotope Laboratories). Glucose-free ImmunoCult™-XF T Cell Expansion Medium (STEMCELL Technologies) was supplemented with 24 mM [U-^13^C]glucose. On day 12 of expansion, CD4+ T cells maintained under CTL:AIM-V or ICM were isolated using human CD4 MicroBeads (Miltenyi) and a MACS LS magnetic column according to the manufacturer’s instructions; CD8+ T cells were collected from the flow through. Separately, the cells were incubated in complete [U-^13^C]glucose CTL:AIM-V or complete [U-^13^C]glucose ICM media supplemented with 300 IU/ml IL-2 (Novartis) for 24 hours at 5% CO2 and 37°C. Metabolic steady state was confirmed on day 12 with 2-NBDG and isotope steady state was confirmed at 24 hours.

To prepare cells for GC-MS analysis, 4 million cells per condition were washed once with ice-cold saline solution (0.9% NaCl solution filtered through a 0.22 μm membrane filter) and resuspended in 1 ml of 50% (vol/vol) methanol (Sigma) (0.22 μm-filtered) cooled at –80°C. Cells were snap frozen at –80°C for 20 minutes and the resulting lysate/methanol mixtures were frozen in liquid nitrogen. Samples were thawed on ice, vortexed, and subject to three freeze-thaw cycles using liquid nitrogen and 37°C water bath. Samples were centrifuged at 10,000 rpm for 10 minutes at 4°C, and then the metabolite-containing supernatant collected. The supernatant was evaporated until dry using a SpeedVac and no heat, and stored at –80°C. Before running GC-MS, norvaline (1 μl) internal standard was added to each sample. Samples were resuspended in 40 μl pyridine containing methoxyamine (10 mg/ml), transferred to GC-MS vials, and heated at 70°C for 15 minutes. 70 μl N-tert-Butyldimethylsilyl-N-methyltrifluoroacetamide (MTBSTFA) was added, samples vortexed, and heated at 70°C for 1 hour before analyzing with GC-MS.

To prepare media for GC-MS analysis, cells were centrifuged and 1 ml of supernatant collected. The samples were frozen at –80°C, thawed on ice, vortexed, and then 25 μl was transferred to a borosilicate tube. Before running GC-MS, 1 μl of norvaline internal standard was added to each sample. Then, 400 μl of methanol, chloroform, and Milli-Q purified water was added to samples. Samples were vortexed and centrifuged at 2,000 rpm for 5 minutes to separate phases. The aqueous phase was collected and evaporated until dry at 42°C using a SpeedVac. The samples were transferred to GC-MS vials and heated at 70°C for 15 minutes before 70 μl MTBSTFA was added. Samples were vortexed and heated at 70°C for 1 hour before analyzing with GC-MS.

Metabolites were analyzed using an Agilent 6970 gas chromatograph and an Agilent 5973 mass selective detector as previously described^25^. GC-MS data was analyzed using Chemstation (Aglient) and Skyline^26^, and graphs were created using GraphPad Prism 8.1.2. The measured distribution of mass isotopomers was corrected for natural abundance of ^13^C^27^. The fractional enrichment of isotopologues were then compared against the fractional enrichment of glucose M+6 for the respective condition. Metabolites with a fractional enrichment of below 0.05 were not considered a significant finding.

Separately, 600 μl media was collected from cells, frozen at –80°C. Glucose and glutamine concentrations in the media were measured using a NOVA BioProfile4 or by colorimetric assay (Biovision).

### Statistical analysis

Statistical analysis was carried out using GraphPad Prism 8.1.2. An unpaired Student’s t-test was used when comparing means of two groups, and one-way ANOVA was used when comparing more than two groups. Differences were considered significant at * p<0.05, ** p<0.01, *** p<0.001, and **** p<0.001.

## Data availability

Processed metabolomics data files are available at https://github.com/vicDRC/metaboData.git. Flow cytometry data will be deposited at flowrepository.

## DECLARATIONS

### Ethics approval and consent to participate

Patient specimens and clinical data were obtained through the BC Cancer Tumour Tissue Repository, certified by the Canadian Tissue Repository Network. All specimens and clinical data were obtained with either informed written consent or a formal waiver of consent under protocols approved by the Research Ethics Board of BC Cancer and the University of British Columbia (H07-00463). Samples are stored in a certified BioBank (BRC-00290).

### Consent for publication

Not applicable.

### Availability of data and material

Processed metabolomics data files will be deposited to github.com. Flow cytometry data will be deposited at flowrepository.

### Authors’ contributions

SM, SK and MK designed and performed experiments, analyzed the data, and wrote the manuscript. JS helped design and performed experiments. JS and RJD ran the mass spectrometry samples and analyzed the raw data. TT and AD irradiated the PBMCs. KSH, JRC, JY, and CS provided reagents and contributed to experimental design. JJL conceived the project and wrote the manuscript.

## LIST OF ABBREVIATIONS

CAR: Chimeric antigen receptor
CD: Cluster of differentiation
CMM: Corning Lymphocyte Serum-Free Medium KBM 581 (Corning)
GC-MS: Gas chromatography-mass spectrometry
ICM: ImmunoCult™-XF T Cell Expansion Medium (STEMCELL Technologies)
IFN: Interferon
IL: Interleukin
HPLM: Human Plasma-like Medium (Dr. Jason Cantor, Gibco)
MFI: Median fluorescence intensity
PBMC: Peripheral blood mononuclear cell
PD-1: Programmed cell death protein 1
PMA: Phorbol 12-myristate 13-acetate
REP: Rapid expansion protocol
ROS: Reactive oxygen species
SEM: Standard error of the mean
TAC: TexMACS™ medium (Miltenyi)
TCA: Tricarboxylic acid
TCR: T cell receptor
T_CM_: Central memory T cell
T_E_: Effector T cell
T_EM_: Effector memory T cell
TIL: Tumor infiltrating lymphocyte
T_N_: Naive T cell
TNF: Tumor necrosis factor
T_reg_: Regulatory T cell

