## Supplementary Figures for "Clinically-relevant T cell expansion protocols activate distinct cellular metabolic programs and phenotypes"

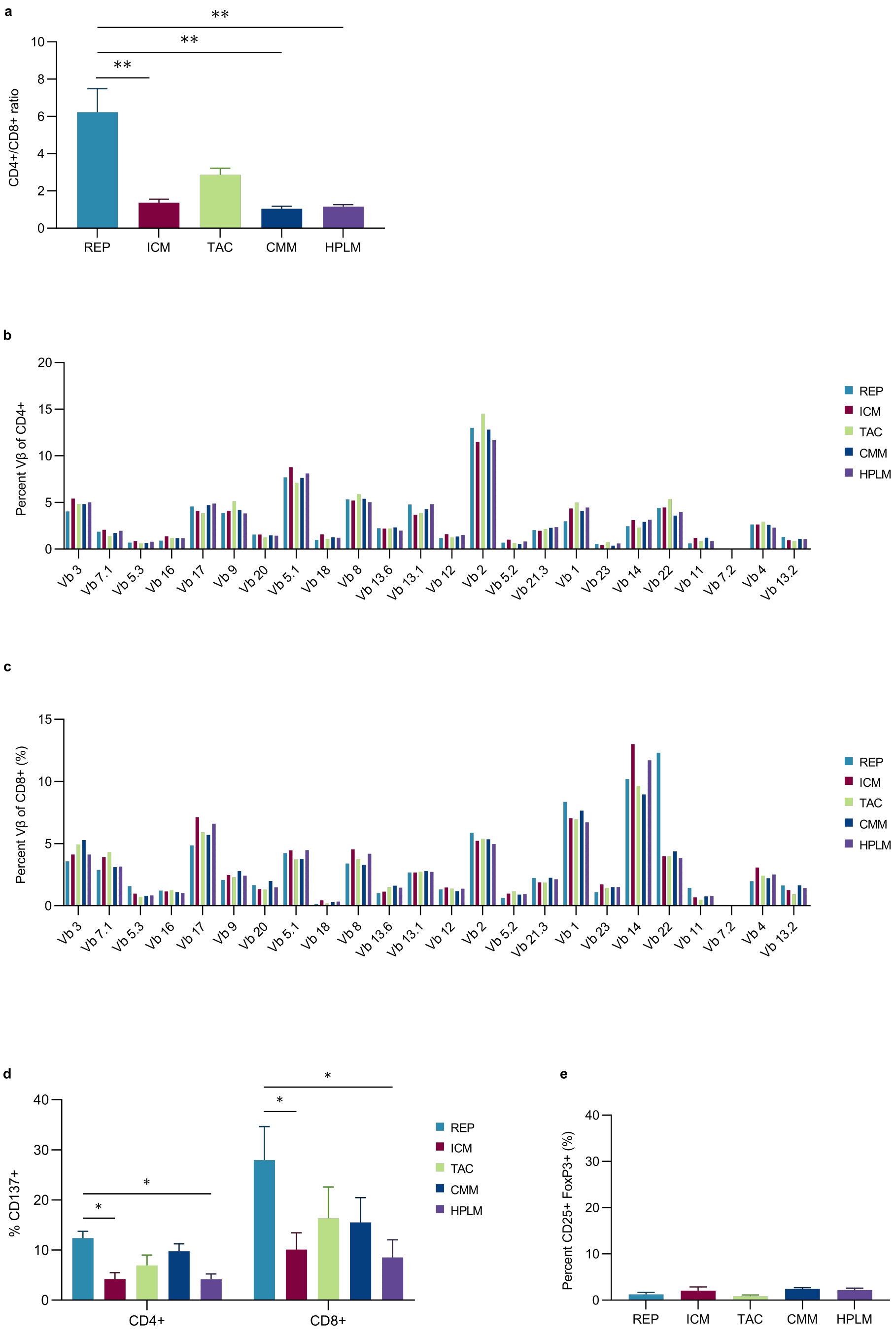


**Supplementary Fig. 1:(a)** Ratio of CD4+ and CD8+ T cells of live CD3+ cells 12 days post-expansion. **(b-c)** Percent of **(a)** CD4+ and **(c)** CD8+ T cells positive for 24 TCR Vβ types following 12 days of expansion. Figure is a representative from one of 3 donors. **(d)** Percentage of CD137 positive CD4+ and CD8+ cells. **(e)** Percentage of $T_{reg}$ cells (CD25+ FoxP3+; gated on live CD4+ cells). (**a**,**d**,**e**) Data are shown as mean of n=3 +SEM from healthy donors. Statistical significance was calculated by one-way ANOVA (* p<0.05, ** p<0.01).

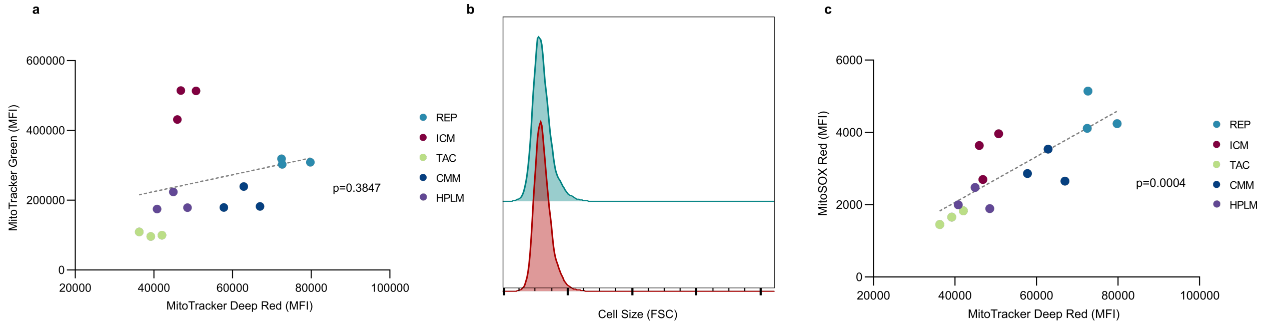


**Supplementary Fig. 2: (a-b)** T cells from three healthy donors were expanded in 5 different conditions for 12 days. **(a)** Correlation between mitochondrial mass (Mitotracker Green) and mitochondrial activity (Mitotracker Deep Red) of live cells. **(b)** Representative plot of cell size of CD3+ T cells in REP (blue) and ICM (red) conditions on day 12. **(c)** Correlation between mitochondrial ROS (MitoSOX) and mitochondrial activity (Mitotracker Deep Red) of live cells. Dashed line represents simple linear regression, p-value determined by Pearson correlation.

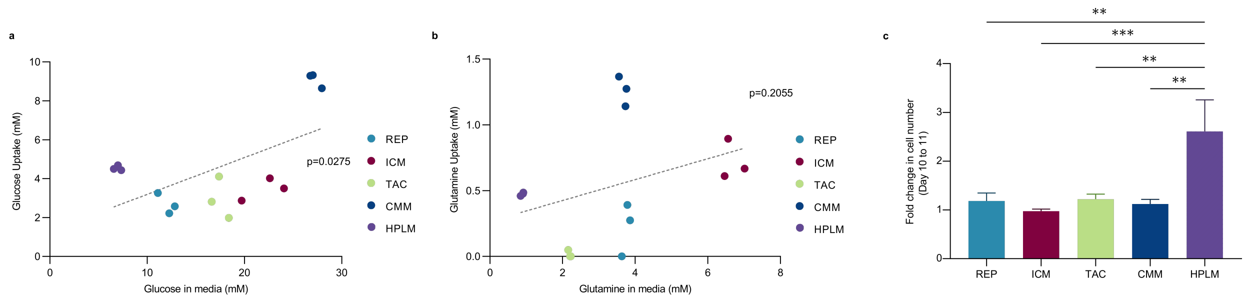


**Supplementary Fig. 3: (a-c)** T cells from three healthy donors were expanded in 5 different conditions for 11 days. Extracellular glucose **(a)** and glutamine **(b)** concentrations in fresh media and spent media after culture for 24 hours in the respective conditions. **(a-b)** Correlation between uptake and concentration in media, dashed line represents simple linear regression, p-value determined by Pearson correlation **(c)** Fold change in cell number from day 10-11 in the respective conditions. **(c)** Data are shown as mean of n=3 +SEM from healthy donors. Statistical significance was calculated one-way ANOVA (** $p<0.01$, *** $p<0.001$).

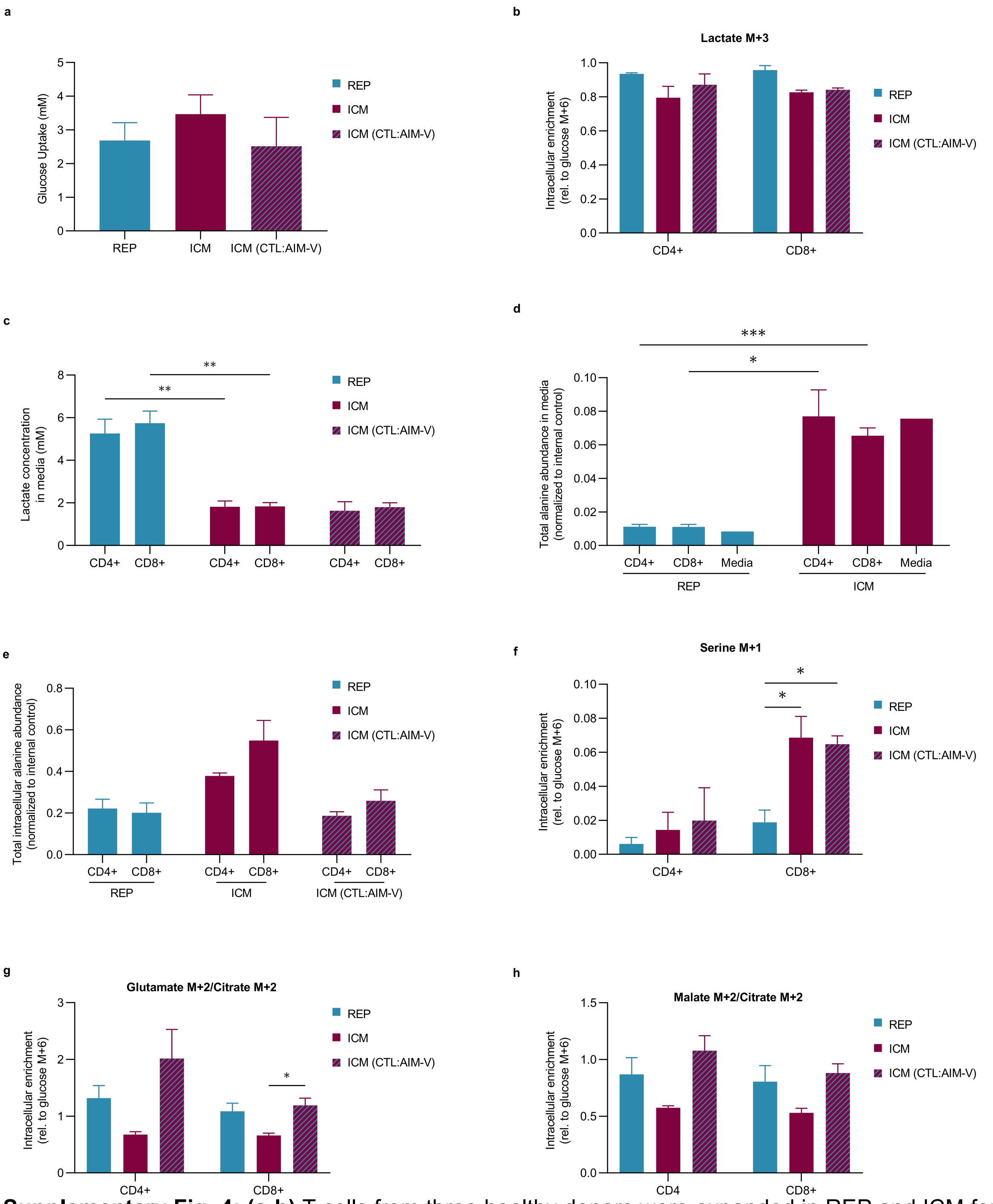


**Supplementary Fig. 4: (a-h)** T cells from three healthy donors were expanded in REP and ICM for 11 days. On day 11, CD4+ and CD8+ cells were separated from the co-culture and incubated in the following [U-$^{13}$C]-glucose conditions for 24 hours: REP cells in CTL:AIM-V (blue bar), ICM cells in ICM (red bar) and ICM cells in CTL:AIM-V (red and blue dashes). **(a)** Glucose uptake from day 11-12 in CD4+ and CD8+ T cells. **(b)** Intracellular lactate M+3 relative to intracellular glucose M+6 enrichment. **(c)** Lactate concentration in media from day 11-12 in CD4+ and CD8+ T cells. **(d-e)** Alanine abundance calculated by totalling alanine isotopologue peak area and normalizing to internal control norvaline peak area. **(f)** Intracellular serine M+1 relative to intracellular glucose M+6 enrichment. **(g)** Glutamate M+2 and **(h)** malate M+2 relative to citrate M+2. Data are shown as mean of n=3 +SEM from healthy donors. Statistical significance was calculated by Student's t-tests (* $p<0.05$, ** $p<0.01$, *** $p<0.001$).

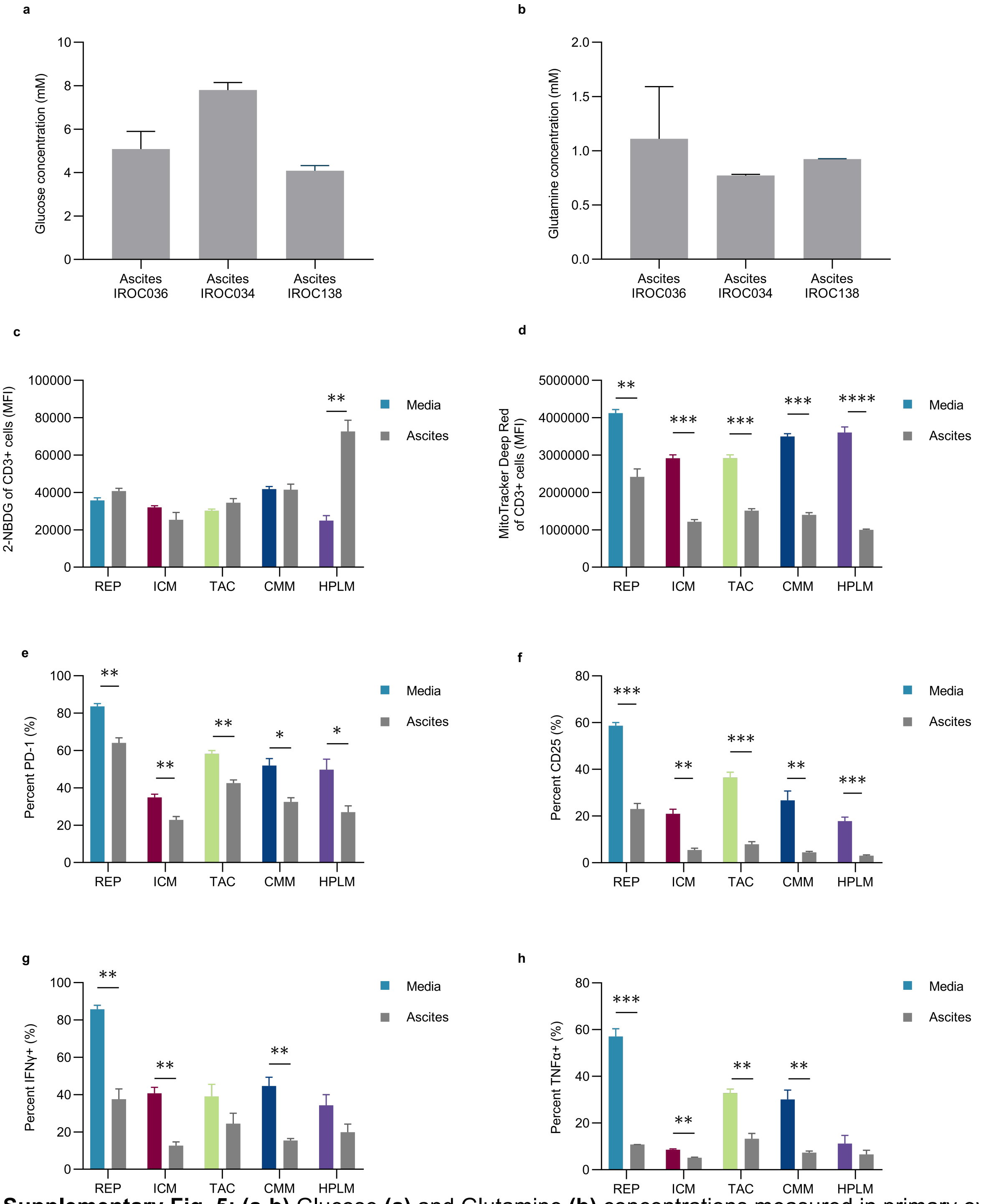


**Supplementary Fig. 5: (a-b)** Glucose **(a)** and Glutamine **(b)** concentrations measured in primary ovarian cancer ascites fluid from three patients. **(c-h)** T cells from three healthy donors were expanded in 5 different conditions over 12 days. On day 12 T cell products were reactivated (CD3/CD28) in media (coloured bars) or ascites (grey bars), T cell metabolism and function was assessed after 2 days. Median fluorescence intensity (MFI) of **(c)** glucose uptake (2-NBDG) and **(d)** mitochondrial activity (MitoTracker Deep Red). Percentage of PD-1 **(e)**, CD25 **(f)**, IFNγ **(g)**, and TNFα **(h)** positive cells in the media and ascites. Data is shown as mean of n=3 +SEM from 3 healthy donors. Statistical significance was calculated by Student's t-tests (* $p<0.05$, ** $p<0.01$, *** $p<0.001$, **** $p<0.0001$).

**Supplementary Table 1:** Flow Cytometry Antibodies

| Fluorochrome | Marker | Expression | Clone | Company | Catalogue number |
|---|---|---|---|---|---|
| FITC | TNFa | Function | MAb11 | BD Biosciences | 554512 |
| PE | IFNy | Function | 4S.B3 | BD Biosciences | 554552 |
| PE CF594 | FoxP3 | Tregs | 236A/E7 | BD Biosciences | 563955 |
| PerCP | CD8 | Effector T cells | RPA-T8 | Biolegend | 301030 |
| PerCP-eFluor710 | CD25 | Activation | 4E3' | Thermo | 46-0257-41 |
| PE-Cy7 | CD45RO | Phenotype | UCHL1 | Thermo | 25-0457-42 |
| --- | MitoTracker Deep Red | Mitchondrial activity | --- | Thermo | M22426 |
| AF700 | CD4 | Helper T cells | RPA-T4 | Biolegend | 300526 |
| APC/Fire750 | CCR7 | Phenotype | G043H7 | Biolegend | 353246 |
| BV605 | CD137 | Activation | 4B4-1 | Biolegend | 309822 |
| BV650 | PD1 | Activation/Exhaustion | EH12.2H7 | Biolegend | 329950 |
| BV750 | CD3 | T cells | SK7 | Biolegend | 344845 |
| --- | MitoTracker Green | Mitchondrial Mass | --- | Thermo | M7514 |
| --- | MitoSOX Red | Mitochondrial ROS | --- | Thermo | M36008 |
| eFlour506 | Viability | Live/dead cells | --- | Thermo | 65-0866-14 |
| --- | 2-NBDG | Glucose uptake | --- | Thermo | N13195 |
| FITC | Vβ | TCR | --- | Beckman | IM3497 |
| PE | Vβ | TCR | --- | Beckman | IM3497 |
| ZombieNIR | Viability | Live/dead cells | --- | Biolegend | 423105 |
